# Colony expansions underlie the evolution of army ant mass raiding

**DOI:** 10.1101/2020.08.20.259614

**Authors:** Vikram Chandra, Asaf Gal, Daniel J. C. Kronauer

## Abstract

Collective behavior emerges from local interactions between group members, and natural selection can fine-tune these interactions to achieve different collective outcomes. However, at least in principle, collective behavior can also evolve via changes in group-level parameters. Here, we show that army ant mass raiding, an iconic collective behavior in which many thousands of ants spontaneously leave the nest to go hunting, has evolved from group raiding, in which a scout directs a much smaller group of ants to a specific target. We describe the structure of group raids in the clonal raider ant, a close relative of army ants. We find that the coarse structure of group raids and mass raids is highly conserved, and that army ants and their relatives likely follow similar behavioral rules, despite the fact that their raids differ strikingly in overall appearance. By experimentally increasing colony size in the clonal raider ant, we show that mass raiding gradually emerges from group raiding without altering individual behavioral rules. This suggests a simple mechanism for the evolution of army ant mass raids, and more generally that scaling effects may provide an alternative mechanism for evolutionary transitions in complex collective behavior.

## MAIN TEXT

Many animal groups, from wildebeest herds to starling murmurations, display complex collective behaviors that emerge from the interactions of individual group members independently following a common set of behavioral rules (Camazine et al., 2001). How these emergent collective behaviors evolve, however, is an open question. One possibility is that natural selection acts on the neural substrate that encodes the underlying behavioral rules. Across species of social insects, for example, workers may respond differently to local cues during nest construction, which could translate into different nest architectures (Mizumoto et al., 2019; Theraulaz and Bonabeau, 1995). Such behavioral rules can evolve rapidly, as has been demonstrated via artificial selection experiments on collective movement in guppies (Kotrschal et al., 2020). In principle, an alternative way to modify collective behavior is to alter group-level parameters, such that the same behavioral rules lead to different collective outcomes. For instance, golden shiners form polarized swarms or milling schools depending on their group size (Tunstrøm et al., 2013). Whether this mechanism is relevant over evolutionary timescales, however, remains unknown. Here we show that army ant mass raiding, one of the most iconic collective phenomena, has evolved from scout-initiated group raiding, and propose that this evolutionary transition in collective behavior was driven substantially by an increase in colony size, rather than changes in the ants’ individual behavior.

Army ants in the subfamily Dorylinae live in huge colonies that contain 104 – 107 workers, depending on the species. They hunt live arthropods, often other ants, in mass raids (Borowiec, 2016; Gotwald, 1995; Kronauer, 2009; Schneirla, 1971) (Table S1). Mass raids begin when workers spontaneously and synchronously leave the nest in “pushing parties” (Leroux, 1977; Schneirla, 1933, 1971). At first, small groups of workers hesitantly leave the nest to explore its immediate vicinity. They lay trail pheromone as they walk, returning after only a few steps out. Ants continue to leave the nest, walking further and further out, confidently following their predecessors’ trail. When they reach untrodden ground, they also hesitate and turn, spreading outwards along the raid front. Over time, this leads to a dynamic fan of ants traveling outwards, leaving a strong, elongating trail back to the nest in its wake (Leroux, 1977; Schneirla, 1933, 1971). In the species with the largest colonies, the ants at the raid front can be so numerous that the raid advances as a swarm (Schneirla, 1971). At the outset, the ants have no information about prey location. However, a few scouts search slightly ahead of the raid front, and when they encounter prey, they lay pheromone trail back to the raid front and recruit nestmates for a collective attack (Chadab and Rettenmeyer, 1975). While army ants themselves have been studied extensively (e.g. (Gotwald, 1995; Kronauer, 2009; Schneirla, 1971)), little is known about their cryptic relatives with much smaller colony sizes. Sporadic and usually partial observations suggest that many non-army ant dorylines conduct scout-initiated group raids, in which scouts find prey before recruiting a raiding party from the nest (Hölldobler, 1982; Hölldobler and Wilson, 1990). It has therefore been suggested that army ant mass raiding might have evolved from scout-initiated group raiding (Gotwald, 1995; Hölldobler and Wilson, 1990; Wheeler, 1918; Wilson, 1958a, 1958b). However, as these species are rarely encountered, no quantitative description of this behavior is available, a formal evolutionary analysis of foraging behavior in dorylines is lacking, and the functional relationship between group raiding and mass raiding is unknown.

We therefore systematically studied foraging behavior in the clonal raider ant, *Ooceraea biroi*, the only non-army ant doryline that can be propagated in the laboratory. In our efforts to establish this species as an experimental model, we have developed high-throughput, automated tracking approaches to monitor individual and collective behavior (Gal et al., 2020; Ulrich et al., 2018). This created the unique opportunity to study doryline foraging behavior quantitatively and under controlled laboratory conditions. In a first experiment, we set up nine colonies each of 25 individually tagged ants, and filmed and tracked their foraging behavior while offering them a single small fire ant pupa once every twelve hours (for experimental details see Materials and Methods). Overall, we analyzed tracking data for 31 raids (Materials and Methods). We found that *O. biroi*, like other non-army ant dorylines, forages in scout-initiated group raids (Movies S1 to S6; for ant foraging terminology see Table S1). We decompose group raids into six distinct phases (Figure 1, A and B; Figure S1). First, in the ‘search’ phase, one or a few scouts explore the arena. Once a scout has discovered food, she examines it briefly before becoming highly excited. In the ‘recruitment’ phase, she runs homeward, and as she enters the nest, the ants inside become active. In the ‘response’ phase, a large proportion of ants inside the nest run towards the scout, exit the nest in single-file, and move towards the food, retracing the scout’s homeward trajectory (Figure 1, A to C). Most ants then stay on or near the food for a few minutes, while some run back and forth between the food and the nest, which we call the ‘pre-retrieval’ phase. Variation in the length of this phase explains most variation in raid length, but its function is currently unknown (Figure 1D, Figure S2). Next, during the ‘retrieval’ phase, one to three ants begin to independently drag or carry the food back home, with no apparent help from their nestmates (Figure 1; Movie S2). Finally, in the ‘post-retrieval’ phase, the last ants outside gradually return to the nest. To visualize the temporal structure of these raids, we aligned and rescaled each phase of each raid, and quantified three informative features: the number of ants outside the nest, the mean distance from the nest, and the sum of the speeds of all ants (Figure 1, E to G). Our analyses show that group raids are highly stereotyped, and mostly vary in the duration of the phases.

**Figure 1:**
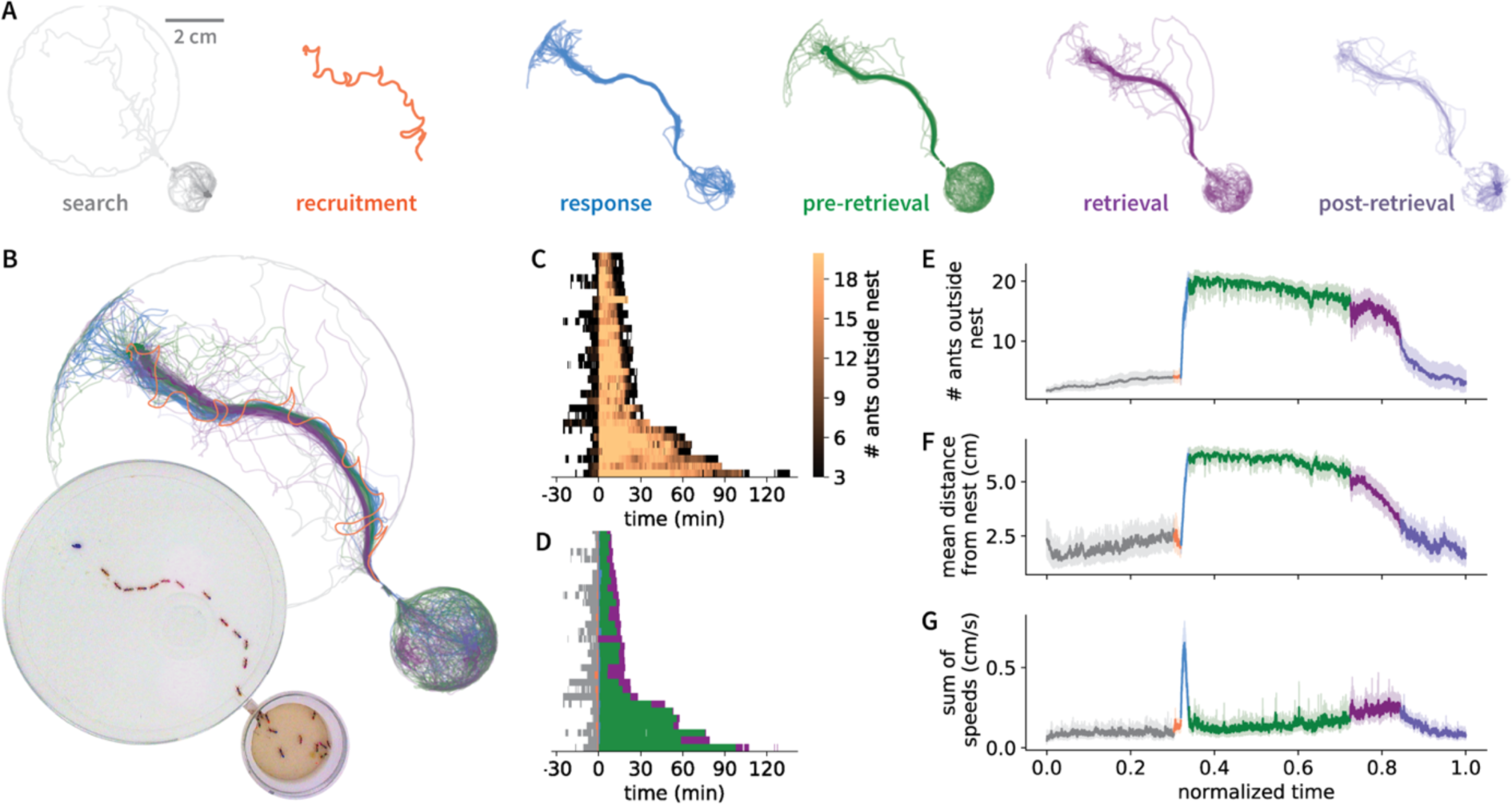
The anatomy of a group raid. (A) Trajectories of ants at each phase of a representative group raid (Movies S1 and S2), separated into six sequential phases (Materials and Methods). The orange track in the ‘recruitment’ phase depicts the path taken by the recruiting ant, whereas tracks in all other phases depict the paths of all ants in the colony. (B) Overlay of trajectories from all six phases. Inset: snapshot of the colony at the peak of the response phase. A short tunnel separates the nest (small circle) from the foraging arena (large circle), and the food (blue spot) is at the top left. (C) Heatmap showing the number of ants outside the nest over time. 31 raids are sorted vertically by their duration and are aligned to the start of recruitment. (D) Representing each phase of each raid by the same color code as in (B) shows that variation in raid length is primarily determined by the length of the pre-retrieval phase (see also Figure S2). We do not show the ‘post-retrieval’ phase here, because it has constant length by definition (Materials and Methods). (E-G) Aligning and rescaling each phase of each raid (Materials and Methods) and plotting the timecourse of the mean number of ants outside the nest (E), their mean distance from the nest (F), and the sum of the speeds of all ants (a measure of collective activity) (G), shows that the temporal structure of group raids is highly stereotyped. The error bands in panels E-G represent the 95% confidence interval of the mean.

To infer the evolutionary relationship between group raiding and mass raiding, we combined our data on *O. biroi* with published descriptions of doryline biology, and mapped relevant life history traits to a consensus phylogeny of the Dorylinae (Table S2) (Borowiec, 2019). Ancestral state reconstructions suggest that the ancestral dorylines lived in small colonies, were specialist predators of ants, and indeed conducted scout-initiated group raids (Figure 2; Figure S3 to S5; Table S3). This supports the hypothesis that army ant mass raiding evolved from group raiding as colony size increased, possibly independently in the New World and Old World army ants (Figure 2) (Hölldobler and Wilson, 1990; Wilson, 1958a, 1958b). It also implies that *O. biroi* might provide mechanistic insight into how these transitions occurred.

**Figure 2:**
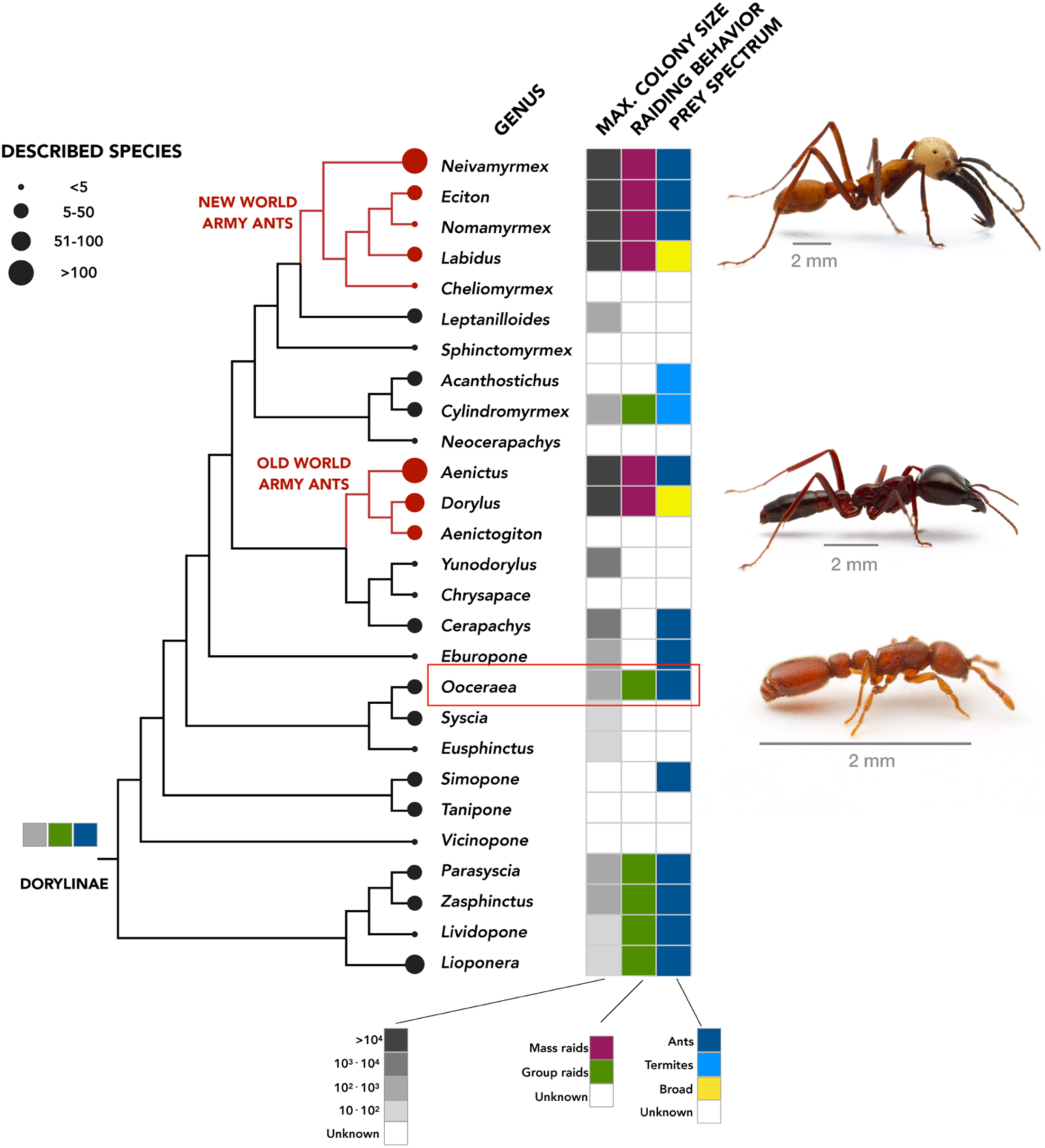
Phylogeny of the Dorylinae, showing all extant genera, along with maximum colony size, type of raiding behavior, and prey spectrum, where known. Ancestral reconstructions on a consensus cladogram (Borowiec, 2019) are shown at the base of the tree (see Figures S3 to S5; Materials and Methods). Photographs from top to bottom show workers of the army ants *Eciton burchellii* and *Dorylus molestus* (photographs © Daniel Kronauer), as well as the clonal raider ant *Ooceraea biroi* (highlighted by a red box; photograph © Alexander Wild).

To understand the evolutionary transition between group and mass raids, it is important to identify homologous phases in the two behavioral sequences. Intuitively, one might compare the response phase of a group raid with the onset of a mass raid, because these are superficially similar: they both represent columns of ants streaming out of the nest. However, homology is best established by identifying the behavioral rules involved in each case. Based on our own observations, as well as previous work on army ants and two distantly related non-army ant dorylines (Chadab and Rettenmeyer, 1975; Gobin et al., 2001; Hölldobler, 1982; Schneirla, 1971), we hypothesized that at least two distinct, scout-derived signals determine the spatial and temporal structure of group raids. First, we asked how the scout activates nestmates. We conducted an experiment in a modified arena that had a porous wall in the middle of the nest chamber, and separate foraging arenas connected to each nest half (Figure 3A). In each trial, food was placed in one foraging arena, and when a scout with access to that arena located the food, she recruited the ants in her nest half, which formed a column that travelled to the food. Shortly after the scout entered the nest, the ants in the other nest half moved towards the wall separating the two halves (Figure 3, A and B; Figure S6; Movie S7). This suggests that the scout releases an attractive recruitment pheromone as she enters the nest, rather than activating nestmates by touch, a contact pheromone, or an undirectional pheromone that signals nestmates to exit the nest chamber without conveying spatial information. Second, we asked whether the scout lays a pheromone trail back to the nest during recruitment, and whether that trail is sufficient to guide the responding ants. Scout-initiated raiding has evolved independently on a few occasions in other ant subfamilies, and in several cases the scout is required to lead the raiding party to the target. In other words, here, information about target location resides primarily in the scout, rather than in a pheromone trail (e.g. (Bayliss and Fielding, 2002; Grasso et al., 1997; Longhurst et al., 1979; Mill, 1984; Topoff et al., 1984)). We found that, in *O. biroi*, the scout usually (in 30/31 raids) does not lead the raiding column (Figure 3C). However, the trajectories of the responding ants closely recapitulate the homebound trajectory of the scout, suggesting that the scout indeed deposits trail pheromone on her way to the nest (Figure 3D). Information about prey location therefore resides exclusively in the scout’s trail. This use of pheromones is highly reminiscent of recruitment at the raid front in army ant mass raids (Chadab and Rettenmeyer, 1975). Together, this suggests that group- and mass-raiding dorylines use chemical information in the same way, and that the recruitment and response phases of a group raid are homologous to recruitment and response at the raid front in mass raids.

**Figure 3:**
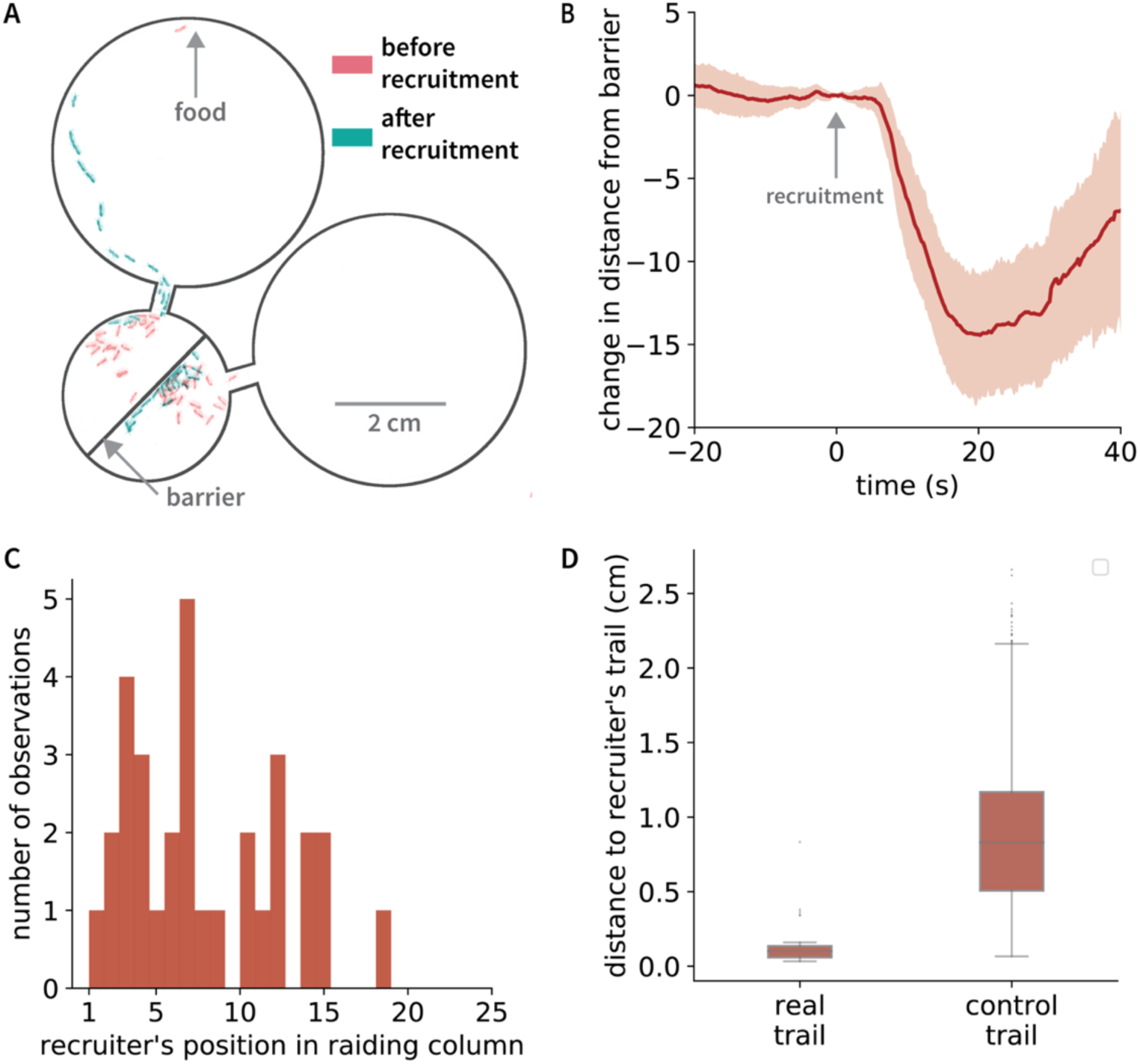
A trail and a recruitment pheromone determine the spatial and temporal structure of group raids, respectively. (A) The recruitment pheromone is attractive and acts at a distance. The image shows a modified nest with a porous barrier down the middle. On the left side, a scout releases recruitment pheromone, causing the ants to leave the nest. The ants on the right side, meanwhile, run towards the barrier instead of leaving the nest. (B) The distance between the barrier and the center of mass of ants on the side opposite to that of the scout as a function of time since recruitment. The center of mass travels towards the barrier after recruitment, which shows that the recruitment pheromone is attractive (n = 31 raids, error band shows 95% CI of the mean). (C) A histogram of the scouts’ position in the raiding column shows that scouts do not typically lead raids. (D) The outbound trajectories of responding ants are significantly closer to their scout’s inbound trajectory than they are to control trajectories of scouts in other group raids, showing that the responding ants indeed follow their scout’s trail to the food (n = 31 raids, Welch’s t-test p<7*10^-29^).

Considering a mass raid to have the same sequence of phases as a group raid, it follows that the onset of a mass raid is actually homologous to the search phase of a group raid (Figure 4A). We therefore asked whether *O. biroi* scouts follow the same basic behavioral rules that translate into spontaneous pushing parties in mass raiding army ants. First, we analyzed our tracking data from colonies of 25 workers to see whether ants incrementally increase their foraging distance by extending previously travelled paths. We found that *O. biroi* often (in 21/31 raids) search an arena that is initially void of trail pheromone in a series of excursions (Materials and Methods). Further analysis of these excursions revealed that, on average, early excursions terminate close to the nest, while later excursions terminate farther away (Figure 4B). Additionally, ants walk faster (Figure S7A) and spend longer outside (Figure S7B) in later excursions and are more likely to follow trail at the beginning, rather than the end, of the outbound leg of each excursion (Figure S7C). This behavior of individual *O. biroi* scouts is highly reminiscent of army ant behavior at the raid front. Taken together, our results suggest that the basic behavioral rules underlying search behavior are conserved between army ants and their non-army ant relatives.

**Figure 4:**
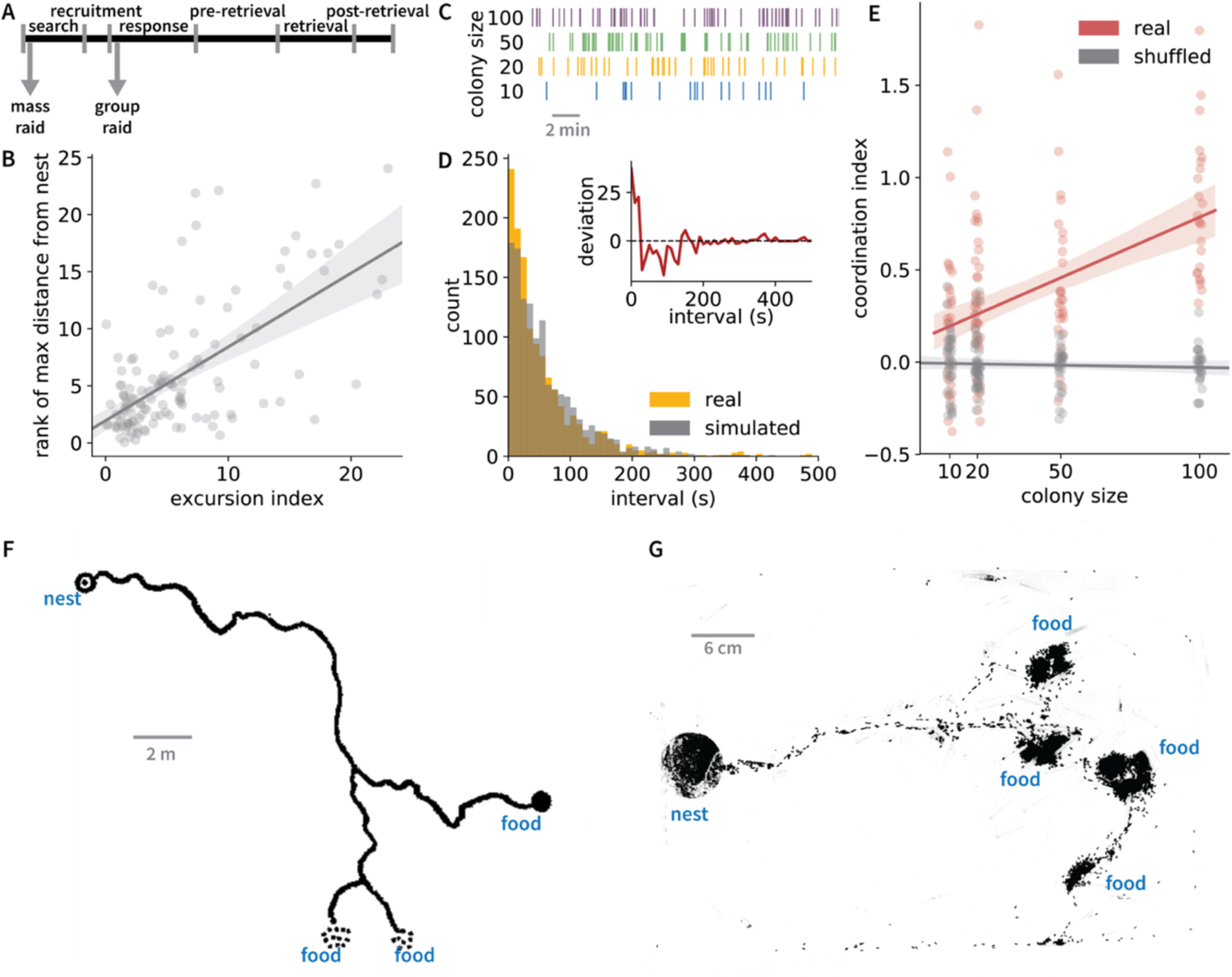
Group raids turn into mass raids with increasing colony size. (A) The onset of a mass raid is homologous to the search phase of a group raid, despite the superficial resemblance to its response phase. Arrows indicate when a column of ants leaves the nest in each type of raid. (B) On average, early excursions terminate closer to the nest than later excursions (n = 127 excursions, colony size 25, linear regression p=4.4*10-16). (C) Four example sequences of nest exit times, sorted by colony size. (D) An example distribution of inter-exit intervals in a colony of size 20. This distribution (in amber) deviates significantly from a simulated exponential distribution (in gray) (Anderson-Darling k-sample test p=0.001). Inset: difference between this distribution and 1,000 simulated exponential distributions, as a function of the interval, showing an increased coefficient of variation (red line is mean difference, with 95% CI). Nest exits in close succession (i.e., short intervals) are overrepresented in the empirical distribution compared to simulated distributions. (E) The coordination index (Materials and Methods) of real inter-exit intervals (red datapoints) increases as a function of colony size (n = 131 exit sequences, linear regression p=4.9*10^-9^), but the coordination index of shuffled interval sequences (gray datapoints) does not (n = 131 exit sequences, linear regression p=0.89). (F) Schematic of a mass raid of the army ant *Aenictus laeviceps*, reformatted with modifications from (Schneirla and Reyes, 1966). (G) Snapshot (background-subtracted and contrast-enhanced; see Materials and Methods) of an *O. biroi* raid in a colony with ca. 5,000 workers. The raid shows all the major features of the army ant mass raid depicted in (F). Error bands in (B) and (E) depict the 95% CI of the regression line.

Despite the similarities in individual behavior, the emergent collective search patterns in a mass raid and a group raid are strikingly different. Unlike in army ants, where workers leave the nest *en masse* to go on a raid, *O. biroi* workers typically leave the nest during the search phase in a seemingly sporadic manner (Figure 1E). To study the temporal structure of search in *O. biroi* and to quantify the synchronicity in the search phase, we conducted an experiment with four colonies of size 20. To control for the possibility that ants behave differently when food is in the arena, we specifically selected periods when the arena was empty (i.e., the ca. 20 hours after each foraging event each day, resulting in a total of 43 distributions). We then recorded each time an ant exited the nest and analyzed the resulting sequences of inter-exit intervals by comparing them to the uncorrelated, exponentially distributed expectation from a random Poisson process (Figure 4, C and D; Materials and Methods). We found that nearly all distributions deviated significantly from the random expectation (Figure S8A), exhibiting increased coefficients of variation (Figure S8B), over-representation of short intervals (Figure S8C), and positive correlations between consecutive intervals (Figure S9A), implying that workers leave the nest in quick succession more often than expected by chance (Figure 4D, Figure S8C). This suggests that, while the apparent synchronicity is weak, a significant positive feedback underlies the search activity of *O. biroi*.

Army ants live in much larger colonies than non-army ant dorylines, and expansions in colony size within the Dorylinae align perfectly with the evolutionary transition to mass raiding behavior (Figure 2). To understand the effect of colony size on the emergent search and raiding behavior, we established *O. biroi* colonies with 10, 50, or 100 workers, alongside the colonies of 20 workers described above. Although these colony sizes do not approach those of army ants, this experiment is nonetheless informative regarding the general scaling effects of colony size. Across all colony sizes, colonies mostly exhibited the same stereotypical raid dynamics (Movie S8), with the number of ants participating in the raids increasing proportionally to colony size (Figure S9B). Analyzing their inter-exit interval distributions (Figure 4C), we observed the same increase in their coefficient of variation compared to the random expectation as before (Figure S8B). Moreover, the correlation between consecutive intervals, as measured by the autocorrelation function of the sequences, markedly increased with colony size (Figure S9A), as did a ‘coordination index’ that we computed from the autocorrelation function (Figure 4E; see Materials and Methods). Thus, as colony size increases, search behavior in *O. biroi* begins to resemble the onset of highly bursty, coordinated army ant mass raids. However, unlike in a full-blown mass raid, these bursts typically attenuate quickly. Nevertheless, we observed multiple events in colonies of ≥50 ants in which positive feedback among the ants spontaneously produced a column that traveled away from the nest, headed by an obvious pushing party that formed without recruitment, in what resembled the onset of a mass raid (Movie S9).

To test whether these scaling effects persist at colony sizes that approach those of army ants, we established two *O. biroi* colonies of roughly 5,000 workers each, an order of magnitude larger than naturally occurring colonies (Tsuji and Yamauchi, 1995), and filmed their raids in large arenas (Materials and Methods). The resulting raids involved thousands of ants and displayed trail bifurcations, simultaneously targeting multiple food sources (Figure 4, F and G; Movie S10; Table S4). Most initial recruitment events now occurred outside the nest and usually at the raid front (43 out of 47). Thus, increasing colony size eventually transforms stereotyped group raids into raids that display all the defining features of army ant mass raids (Table S1).

Together, our results suggest that all doryline ants share fundamental rules of search and recruitment behavior. At small colony sizes, these rules manifest as scout-initiated group raids. However, as colony size increases, either within species or between species across evolutionary time, these rules gradually give rise to spontaneously initiated mass raids in which many ants leave the nest in quick succession, advance in pushing parties, and recruit at the raid front rather than at the nest. The difference between search behavior in group raiders and mass raiders may thus be largely driven by the effects of increasing colony size. Although the mechanism underlying this change in group dynamics is currently unknown, the transition from an irregular, low-activity state to a synchronized, high-activity state with increasing colony size or density is reminiscent of similar transitions observed in other complex systems, including excitable membranes, neural networks, cooperative microorganisms, and locust swarms (Buhl et al., 2006; Gregor et al., 2010; Hodgkin and Huxley, 1952; Schneidman et al., 2006). This dynamical transition underlies the emergence of army ant mass raiding and constitutes a striking example of an evolutionary change in collective behavior that need not require modification of neural circuitry.

## ACKNOWLEDGEMENTS

We thank Leonora Olivos-Cisneros and Stephany Valdés Rodríguez for assistance with ant maintenance, Jim Petrillo and the Rockefeller University Precision Instrumentation Technologies facility for assistance with arena construction, the Rockefeller University High Performance Computing Core for computing infrastructure, and the Kronauer lab for helpful discussion. This work was supported by a Faculty Scholars Award from the Howard Hughes Medical Institute and the National Institute of General Medical Sciences of the National Institutes of Health under Award Number R35GM127007, both to D.J.C.K. The content is solely the responsibility of the authors and does not necessarily represent the official views of the National Institutes of Health. A.G. was supported by the Human Frontiers Science Program (LT001049/2015). This is Clonal Raider Ant Project paper #16.

## AUTHOR CONTRIBUTIONS

V.C., A.G., and D.J.C.K. designed the study; A.G. designed and built the tracking setup, and wrote the tracking software; V.C. and A.G. designed and built the raiding arenas; V.C. conducted the experiments and evolutionary analyses; V.C. and A.G. analyzed the tracking data; V.C. and D.J.C.K. wrote the manuscript with feedback from A.G.; D.J.C.K. supervised the project.

## MATERIALS AND METHODS

### Colony maintenance

*Ooceraea biroi* colonies were maintained in the lab at 25°C in boxes with a water-saturated plaster of Paris floor. Like many other doryline ants, colonies of this species undergo stereotypical cycles, alternating between reproductive phases, during which the ants lay eggs and do not forage, and brood care phases, during which colonies contain larvae, and workers forage for food. During the brood care phase, experimental colonies were fed with frozen *Solenopsis invicta* brood. All experiments were performed using ants from clonal line B (Kronauer et al., 2012). For all experiments other than the one with very large colonies, all ants were one month old, were from the same source colony, and had been reared under the same conditions.

### Behavioral tracking setup

Behavioral experiments were conducted in artificial arenas constructed from layers of cast acrylic, with a plaster of Paris floor. Each arena was a square of side 10 cm, in which we laser-cut a nest chamber and a foraging arena, connected to each other by a narrow tunnel (see Figure 1). The nest chamber had a diameter of 2 cm, the tunnel was ~2 mm wide and ~6 mm long, and the foraging arena had a diameter of 6.5 cm. The floor of the foraging arena was covered with vapor-permeable Tyvek paper to make it less attractive as a nesting site and discourage colonies from emigrating there, while keeping it suitable as a foraging arena. For all experiments in these artificial arenas, ants were introduced to the nest chamber at the start of the reproductive phase. During this period, the tunnel was sealed to prevent ants from entering the foraging arena. 2-4 days after introduction, the ants laid eggs in the nest chamber. Ten days later, the eggs hatched into larvae. 4-6 days after this, when the larvae were in their third or fourth instar, we placed food (i.e., a single frozen *S. invicta* pupa) in the foraging arena, unsealed the tunnel, and filmed the ants foraging.

We filmed colonies at 5-10 Hz and 2592×1944 pixel resolution, using webcams (Logitech C910) in enclosed containers with controlled LED lighting at ~27°C and ~60% humidity.

### Tagged-ant experiment

Nine colonies of ants were established from a single cohort of one-month old ants that were entering the reproductive phase. Each colony consisted of 25 ants, and each ant was tagged with an ordered pair of color dots that was unique within the colony. Specifically, each ant was painted on her thorax and gaster with one of five colors of oil-paint markers (uni Paint Markers PX-20 and PX-21), a previously-used technique (Gal et al., 2020; Trible et al., 2017; Ulrich et al., 2018). At the end of the experiment, we counted all larvae, and found that each colony had between 20 and 25 larvae. In other words, the larvae:adults ratio (a known source of variation in colony foraging – see (Ulrich et al., 2016)) was close to 1:1 in all colonies.

For the eight days of the tracking period (i.e., when the larvae were between ~5-13 days old), every 12 hours, we cleaned each foraging arena with water (to remove trail pheromone from the previous foraging event), and placed a single *S. invicta* pupa (infused with 0.05% bromophenol blue to aid visualization – Movie S1; Figure 1B) at its far end. We then unsealed the tunnel and allowed the ants to explore the arena. We filmed the arena for roughly four hours thereafter, at 10 frames per second (fps), after which we resealed the arena. For the first five days (i.e., the first ten foraging events), each colony was given a small (worker-destined) *S. invicta* pupa. For the next three days, we presented colonies with large (queen-destined) or small (worker-destined) pupae in alternation. The difference in feeding did not affect the coarse structure of the colonies’ foraging behavior. Here, we do not differentiate between these foraging events, and we will analyze the fine differences between them in a subsequent publication. In some cases, colonies emigrated to the foraging arena. For the next event in such colonies, if the ants had not moved back to the nest chamber, we presented them with a *S. invicta* pupa but did not record foraging. All chambers had their plaster floor watered periodically to saturation.

In sum, we recorded 90 foraging events (i.e., events in which the ants were provided food, discovered it, and the food entered the nest) across nine colonies. Of these, 22 events ended in emigration. In 18 events, the ants appeared to eat the *S. invicta* pupa *in situ* (although we cannot exclude the possibility that they tore it into small pieces before carrying it home, and we cannot be certain that only adults ate the food). The 50 remaining events ended in retrieval – i.e., with the ants transporting the pupa into the nest. We never observed emigration again in subsequent experiments, and only observed a single further instance of eating *in situ*, possibly due to subtle differences in experimental design. Thus, we excluded these events from our analysis here. Of the 50 events that ended in retrieval, 19 were excluded from analysis due to failures in data acquisition or cases where the colony was unsettled at the time of food presentation. Our final dataset thus consisted of 31 foraging events from seven colonies.

### Annotation of group raid phases

Based on our manual observations of the raids, we identified six discrete, sequential phases of each raid. We defined the ‘search’ phase as the period beginning at the start of the video and ending at the time at which the next phase (i.e., ‘recruitment’) begins. For the group raids that we analyze here, scout ants necessarily located the food during the search phase. The recruitment phase begins when a scout leaves the food and runs homeward, and it ends when the scout recruits her nestmates, which commences the ‘response’ phase. The recruitment phase only includes successful recruitment. In some cases, scout ants may run homeward from the food without initiating a response; however, as we cannot judge whether these instances constitute attempted recruitment, we do not use them to define the beginning and end of the recruitment phase. We define the beginning of the response phase as the first ant of a column leaving the nest, and the end as the moment when the tail of the column reaches the food. This commences the ‘pre-retrieval’ phase, which ends when ants begin to move the food back home. We define the ‘retrieval’ phase as beginning when the position of the food has noticeably changed and ending when the food enters the nest. We define the final phase, ‘post-retrieval’, as beginning when the food has entered the nest and arbitrarily end it 500 seconds later.

For all raids, we manually annotated the corresponding videos, specifically recording five timepoints that allow us to define these six phases. These timepoints are the time at which a scout leaves the food on her recruitment run, the time at which the leader of the column of ants responding to recruitment enters the foraging arena, the time at which the last ant in the column arrives at the food, the time at which the position of the food begins to change, and the time at which the food enters the nest. In colonies of 25 ants, these timepoints may be recorded with minimal subjectivity, as assessed by repeated annotations of the same raids, and by comparisons of recorded timepoints between observers (data not shown). For all raids analyzed here, a single observer (V.C.) annotated all videos. We also recorded the identities of the scouts that successfully initiated raids, and all ants that contributed to retrieving food.

### Tracking tagged ants

Videos from this experiment were processed using anTraX (Gal et al., 2020) to produce the spatial (xy) coordinates of each ant in the colony during each of the events. Tracking quality was quantified using the assignment error (see (Gal et al., 2020) for definition) for each colony and each event separately. As in the context of this paper we have used individual trajectories for the analysis of the recruitment and response phases, we have quantified the assignment error in the time period starting just before start of recruitment phase and ending just after the end of the response phase. The average tracking error was estimated to be 1.8%, with the largest error across all events being 5%.

### Visualization of the average raid structure

While the temporal ordering of the phases is identical across raids, the duration of each of the phases vary considerably between events (Figure 1D, Figure S2). In order to analyze the average time course of colony activity during the raid, we computed the mean duration of each of the phases across all 31 raiding events. We then rescaled each phase of each raid so that it equaled the mean phase duration. Rescaling was done by dividing the timepoints of each phase in each event by the ratio between the average phase duration and the current phase duration. We then interpolated and resampled each of the computed measures (number of ants outside the nest, their average distance from the nest, and the sum of their absolute velocities), so that all events had the same time axes, and applied a moving average filter with a window size of 1 second to smooth out tracking noise. The average of these rescaled time-dependent measures, together with their 95% confidence intervals is shown in Figure 1, E to G.

### Analysis of the scout’s position in the raiding column

To ask whether the scout led the raid, we ranked her position in the raiding column in each raid. To do this, we took advantage of the fact that in all analyzed raids, the responding ants walked in a single file. We ranked all ants by the time they crossed the halfway mark between the nest and the food (Figure 3C). Observations of the videos suggested that changes in the ants’ ranks were minimal (i.e., they did not often overtake each other), and selecting alternative points at which to rank the ants did not noticeably alter the distribution of the scout’s rank across raiding events (data not shown).

### Analysis of trail following during the response phase

To ask whether the ants in the response phase follow the specific trail laid by the scout in the recruitment phase, we asked whether the xy coordinates during their outbound journey were closer to the xy coordinates of the recruiting scout during her inbound journey than expected by chance.

For each raid, let the set of the recruiter’s xy coordinates be:

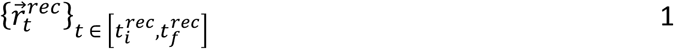

where 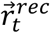 represents the xy coordinates of the recruiter at time *t*, 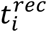 is the time at the start of the recruitment phase, and 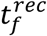 is the time at the end of the recruitment phase.

Similarly, the set of all xy coordinates of all responding ants is:

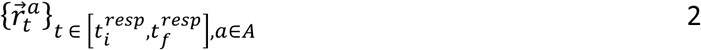

where 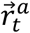 represents the xy coordinates of ant *a* at time *t*, 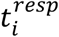 is the time at the start of the response phase, 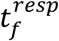 is the time at the end of the respons phase, and A is the set of ants that participate in the response to recruitment.

For each timepoint in the response, and for each ant participating in the response, we define 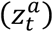 as its minimum distance to the recruiter’s track:

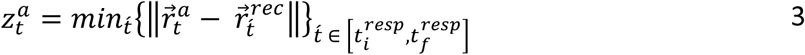

For each raid event, we then computed a measure of trail following, defined as:

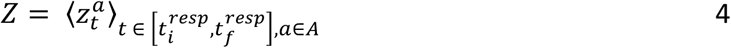

If the ants are not following the recruiter’s trail, we might still expect *Z* to have a relatively low value, because the positions of the nest and the food remain constant across each raid (and thus substantially constrain the initial and final xy coordinates of each ant’s trajectory). To account for this inherent spatial structure in our null expectation, we compared the set of response xy coordinates to the xy coordinates of scouts from all raids other than their own. For each set of response coordinates, we thus generated 30 minimum-distance values. We then compared the distribution of 31*30 ‘control’ *Z* values to the distribution of 31 true *Z* values using a Welch’s t-test.

### Detection and analysis of excursions in the search phase

We define an excursion as the trajectory of a scout from the moment she is leaving the nest, to the moment she enters back. To identify excursions in our data, for each ant, we identified all pairs of transitions across the nest threshold and extracted the trajectory segments between them. For each excursion, we then calculated a number of summary features: its duration, its maximum distance from the nest, and the ant’s mean speed. We excluded excursions in which ants traveled >= 3.5 times their maximum distance from the nest, as these represent cases where the scout travels along the arena’s wall or is walking in circles in the arena. We then ranked these values within each event and plotted the excursion rank versus its index in the event across all events (Figure 4C, Figure S7, A and B).

To ask how ants follow trails during these excursions, we also selected the outbound leg of each excursion by truncating the excursion at the time at which the ant reached its maximum distance (in that excursion) from the nest. For each xy coordinate in each outbound leg, we classified it as either on-or off-trail, depending on whether it mapped to a previously occupied pixel on a 100×100 pixel binary map (where each pixel represents a square of side 1mm) of all previous ant locations, excluding the focal excursion, in that search phase. We then rescaled all such binary sequences to be the same length so that we could align the beginning and end of the outbound legs of each excursion (Figure S7C).

### Barrier experiment

To study the nature of recruitment, we modified our artificial arenas as depicted in Figure 3. We laser-cut cast acrylic porous barriers of 0.8 mm thickness, with multiple holes with a diameter of ~50 µm, so that ants could not contact each other from across such barriers but could communicate via volatile pheromones. Each barrier was placed in the middle of a nest that had two foraging arenas, essentially creating two nests separated by this porous barrier. We established colonies of 20 one month old, phase- and genotype-matched ants in each nest half in each of eight replicate nests. The ants laid eggs in each nest half two days later. In the subsequent brood care phase, each day (except for a handful of days interspersed through the experiment when we fed and watered all colonies while preventing them from leaving their nest halves), we placed a single *S. invicta* pupa in the foraging chamber of one nest half of each artificial arena, alternating which half received food each day. In this experiment, a number of colonies often failed to detect the food (because the ants never left their nest). Nonetheless, we recorded 35 instances of foraging in five artificial nests across a two-week period. Of our 35 replicate events, we excluded four events from a single colony from further analysis, because the scout in these events did not enter the nest, or because (in one case) the colony was too active in the search phase for effective recruitment.

As tracking the ants in the dense chamber is impractical, we used an alternative approach to understand the recruitment dynamics in the nest. We subtracted each frame in the video from the background image (an image which includes all image features, but without the ants; see (Gal et al., 2020) for the procedure used to generate these images), and converted it to gray scale. As ants are darker than the background, the value of each pixel, *g*_*i*_, in this image was taken as the probability that it contains an ant. We then compute the center of mass coordinates for each “half-colony” by summing over all pixels that belong to it:

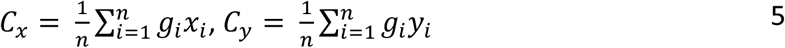

where C refers to the centroid’s coordinates, *x* and *y* refer to the coordinates of each pixel, and *g* refers to the pixel gray-value.

For each frame, we found the position of the centroid, and recorded its distance to the barrier separating the two nest halves as the length of the perpendicular from the centroid to the barrier. We then aligned the time series of centroid distance (from the barrier) to the time of recruitment (which we define as the time at which a majority of ants on the scout’s nest half are activated and begin to move), and averaged across events (Figure 3B, Figure S6).

For the statistical analysis comparing distances before and after the scout releases recruitment pheromone (Figure S6A), we manually selected a frame from each video roughly 1-2 seconds before release, and compared the distance of the centroid from the barrier at this timepoint to its distance 20 sec later. To ensure that our manual selection of the initial frame was accurately identifying a time shortly before recruitment, we also defined the “ant mass” *M* in each nest half by:

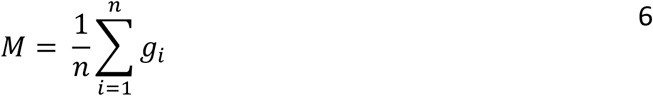

We then plotted the ant mass in the scout’s nest half over time from the time recruitment, and found that shortly after the initial frame, this ‘ant mass’ decreased sharply – an indication that the ants in the scout’s nest half actually left the nest in response to recruitment (Figure S6).

### Colony size experiment

To ask how increasing colony size altered the structure of search behavior, we established 3-4 colonies each of 10, 20, 50, or 100 untagged workers. As before, all workers were one month old, and were selected from a single cohort from a large source colony. They were placed in artificial arenas identical to those used in the tagged-ants experiment when they were entering the reproductive phase, and laid eggs simultaneously in their new nests shortly thereafter. In the subsequent brood care phase, when their larvae were ~5 days old, we began tracking. Here, every day for 10 days, we gently transferred ants in the foraging arena into the nest, sealed the connecting tunnel, cleaned the foraging arena with water, saturated the plaster base of each colony, and placed food (a single small *S. invicta* pupa) in the foraging arena before reopening the tunnel and starting tracking. Roughly four hours later, we then fed each colony in proportion to their colony size (to control their nutritional states). Specifically, we placed *S. invicta* pupae inside each nest, maintaining a constant 1:10 food items:ants ratio. On rare occasions when a colony did not locate the food in the arena within four hours, we placed it inside the nest. We then continued filming the colony for the next ~20 hours. We repeated this process through the brood care phase, until the larvae had pupated. This experimental design allowed us to study how varying colony size alters the structure of the raid (Figure S9B), and more importantly, how it alters the behavior of ants searching for food when there is no food in the arena – the primary focus of our statistical analyses (Figure 4).

### Exit counting analysis and controls

To analyze the temporal structure of search behavior, we recorded the time at which each ant exited the nest (and entered the foraging arena). We used anTraX to track ant movement while in the foraging arena. Since the ants were not individually tagged in this experiment, we did not obtain complete trajectories, but rather a collection of short tracklets, some of which were single-ant and some were multi-ant (Gal et al., 2020). We marked the entrance to the tunnel and filtered all tracklets that originated with an ant emerging from the tunnel (all tracklets that have their first blob overlapping with the entrance mark and have no parent tracklets, or multi-ant tracklets with only one single-ant tracklet parent that start at the tunnel entrance). For each of these tracklets we recorded the first frame as an “exit time” of one ant. While the false positive rate of this detection process is minimal, the false negative (unrecorded exits) is more substantial, as some cases where ants leave the nest in close proximity, which prevents their segmentation, are recorded as single exits. However, for all the analyses described below, these errors work to decrease the reported effect.

Overall, across all colony sizes, we had 150 timeseries of intervals between subsequent nest exits. We excluded samples (i.e., timeseries) that had fewer than 200 total exits from subsequent analysis. As our analysis was focused on short-term activity fluctuation, we detrended each timeseries with third-degree polynomials to account for slow modulations of activity that might correspond to effects such as buildup of colony hunger, circadian cycles, etc. For each timeseries, we then assessed the autocorrelation for the first ten lags of the autocorrelation function (Figure S9A). The mean autocorrelation was higher for larger colony sizes at most initial lags. To quantify a ‘coordination index’ *C* for ants leaving the nest together, we summed the unbiased autocorrelation over the first ten lags, and compared this value across samples:

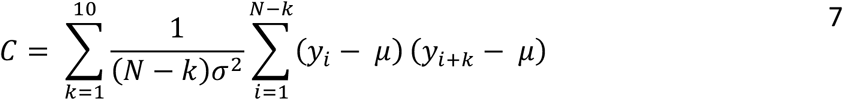

where *Y*_*i*_ refers to the i-th inter exit interval in the detrended sequence,

*k* refers to the lag,

*μ* refers to the empirical mean,

*σ* refers to the empirical standard deviation,

and *N* refers to the size of the inter-exit sequence.

### Quantifying the number of ants that participate in the raid

As a proxy for the true number of ants involved in raids, we used the maximum number of detected blobs outside the nest in any single frame throughout the raid, whether these blobs corresponded to individual or several ants. While this is an underestimation of the real number of ants participating in the raid, it provides an estimate that is sufficient for the purpose of testing whether larger colonies have more total participation (Figure S9B). Moreover, the negative bias in the estimation increases as a function of colony size (data not shown).

### Enlarged *O. biroi* colony experiment

We established two *O. biroi* colonies in the brood care phase with roughly 5,000 workers each. All workers in both colonies were of clonal line B and included multiple age-classes, representative of natural colonies. Preliminary experiments suggested that colonies of this size settle relatively rapidly, and we found that after 12 hours in a new nest, the colonies behaved qualitatively indistinguishably from colonies that had lived in a nest for arbitrarily long periods. For each foraging event, we anesthetized each colony with CO_2_ and transferred it into a new arena (roughly 60cm × 34cm) with a fresh plaster of Paris base and a circular nest chamber (radius 6cm) with a single sealed exit.

*O. biroi* workers have a strong thigmotactic tendency, and in large, featureless arenas, they spend substantial proportions of time following the outer walls. To ameliorate this effect, we scattered a number of small, transparent acrylic bricks (3cm × 0.3cm × 0.3cm) throughout the arena. Pilot experiments suggested that introducing these bricks inside the arena would enable the workers to follow the short local edges, diminishing the amount of time they take to locate the food and creating more naturalistic conditions. Additional pilot experiments showed that adding such edges or changing arena size did not qualitatively affect the ants’ ability to raid.

Roughly 12-16 hours after introducing each large colony to its new nest, we placed 3-7 piles of fire ant brood far from the nest, and then unsealed the nest exit and allowed each colony to explore the arena. We filmed each colony’s foraging behavior for the next ~24 hours. We repeated this process seven times for one colony and four times for the other, with 1-3 days between subsequent foraging events. Together, we filmed eleven foraging events in the brood care phase in these large arenas, of which we excluded one because the ants were alarmed at the start of filming. We manually annotated the remaining foraging events to assess whether recruitment occurred inside or outside the nest, whether or not recruitment events resulted in bifurcation of the trail, and to estimate approximately how many ants participated in the raid (Table S4).

To create the image shown in Figure 4G, we selected a representative snapshot from the middle of one of these raids, performed background-subtraction and then uniformly increased the contrast of the image to make the ants more visible.

### Ancestral state reconstructions

We used the phylogenetic consensus topology of the Dorylinae from (Borowiec, 2019). We searched the natural history literature on doryline ants to find information on character states for a number of characters: colony size, prey spectrum, and various features of foraging behavior (raid initiation, recruitment, number of ants in the raid, and trail bifurcation) that are characteristic of either group or mass raiding behavior (Tables S1 and S2). Since there is very little evidence from multiple species within each genus (and little quantitative data anywhere in the Dorylinae), we chose to collapse character states for each trait into a genus-level categorical assessment. There were no major ambiguities within any genus.

To infer the ancestral states of foraging behavior (Figure S5), we classified each genus as either a group raider, a mass raider, or as ‘unknown’, based on their four foraging characters’ states (Table S2). There were no inconsistencies across the four characters for any genus - i.e., any species with one character state typical of group raiding had other character states also typical of group raiding, or had no information regarding other character states. Thus, if a genus had at least two known character states, we classified it as either a group or mass raider. We classified genera with information for one or no characters as ‘unknown’.

We then reconstructed ancestral states for maximum colony size (Figure S3), prey spectrum (Figure S4), and raiding behavior (Figure S5) using maximum parsimony (MP) and maximum likelihood (ML) with a one-parameter Markov k-state model, both implemented in Mesquite (Maddison and Maddison, 2019) (Table S3). Given the paucity of character data, we interpret this reconstruction largely qualitatively, ignoring inferred character states for all intermediate nodes except the doryline most recent common ancestor (MRCA).

## Notes

### Competing Interest Statement

The authors have declared no competing interest.

