## Supplementary tables, figures, and movie legends for "Colony expansions underlie the evolution of army ant mass raiding"

| Type | Raid initiation | Initial site of recruitment | Number of ants participating in raid | Trail bifurcations |
|---|---|---|---|---|
| **Group raid** | Scout-initiated | Inside nest | Dozens to hundreds | No |
| **Mass raid** | Spontaneous* | Outside nest (at raid front)** | Thousands to millions | Yes |

Table S1: Comparison between the two different types of known foraging behavior in the Dorylinae. Following historical precedent (Berghoff, 2002; Wilson, 1958b; Witte, 2001), we use the terms 'group raid' and 'mass raid' to distinguish two syndromes of raiding behavior. The table identifies four distinguishing features of each type. Although an ant colony could in principle have a combination of 'group raid' and 'mass raid' features, in practice such colonies have not been observed in nature (see Table S3).

(*) Initiation of mass raids is 'spontaneous' with respect to the discovery of prey. In surface-active species, mass raids are sometimes triggered by sunrise or follow an apparently circadian rhythm (Rettenmeyer, 1963; Schneirla, 1971; Topoff et al., 1980).

(**) recruitment outside the nest may be followed by further recruitment inside the nest– this classification is hierarchical.

| GENUS | # described species | max. colony size (o.m.) | army ant? | diet | raid initiation | recruitment | # ants in raid | trail bifurcation | raiding behavior | references |
|---|---|---|---|---|---|---|---|---|---|---|
| *Neivamyrmex* | 130 | 100000 | Yes | ants | spontaneous | outside nest | many | yes | mass | (Borowiec, 2016; Rettenmeyer, 1963; Topoff et al., 1980) |
| *Eciton* | 29 | 1000000 | Yes | ants | spontaneous | outside nest | many | yes | mass | (Schneirla, 1933, 1934) |
| *Nomamyrmex* | 2 | 1000000 | Yes | ants | ? | ? | many | yes | mass | (Borowiec, 2016; Rettenmeyer, 1963) |
| *Labidus* | 9 | 1000000 | Yes | generalist | ? | ? | many | yes | mass | (Borowiec, 2016; Monteiro et al., 2008; Rettenmeyer, 1963) |
| *Cheliomyrmex* | 4 | ? | Yes | ? | ? | ? | ? | ? | ? | (Gotwald, 1971; O'Donnell et al., 2005; Wheeler, 1921) |
| *Leptanilloides* | 19 | 100 | No | ? | ? | ? | few | ? | ? | (Brandão et al., 1999; Donoso et al., 2006) |
| *Sphinctomyrmex* | 3 | ? | No | ? | ? | ? | ? | ? | ? | |
| *Acanthostichus* | 23 | ? | No | termites | ? | ? | ? | ? | ? | (Brown, 1975; MacKay, 1996) |
| *Cylindromyrmex* | 10 | 100 | No | termites | scout-initiated | inside nest | few | no | group | (Gobin et al., 2001; Mariano et al., 2004) |
| *Neocerapachys* | 2 | ? | No | ? | ? | ? | ? | ? | ? | |
| *Aenictus* | 183 | 100000 | Yes | ants | spontaneous | ? | many | yes | mass | (Borowiec, 2016; Chapman, 1964; Schneirla and Reyes, 1966) |
| *Dorylus* | 60 | 10000000 | Yes | generalist | spontaneous | outside nest (inferred) | many | yes | mass | (Berghoff, 2002; Berghoff et al., 2002; Borowiec, 2016) |
| *Aenictogiton* | 7 | ? | Yes* | ? | ? | ? | ? | ? | ? | (Borowiec, 2016) |

| | | | | | | | | | | |
|---|---|---|---|---|---|---|---|---|---|---|
| *Yunodorylus* | 4 | 1000 | No | ? | ? | ? | few | ? | ? | (Borowiec, 2016) |
| *Chrysapace* | 3 | ? | No | ? | ? | ? | ? | ? | ? | (Borowiec, 2016) |
| *Cerapachys* | 5 | 1000 | No | ? | ? | ? | few | ? | ? | (Borowiec, 2016) |
| *Eburopone* | 1 | 100 | No | ants | ? | ? | ? | ? | ? | (Borowiec, 2016) |
| *Ooceraea* | 13 | 100 | No | ants | scout-initiated | inside nest | few | no | group | (Borowiec, 2016) |
| *Syscia* | 5 | 10 | No | ? | ? | ? | ? | ? | ? | (Borowiec, 2016) |
| *Eusphinctus* | 2 | 10 | No | ? | ? | ? | ? | ? | ? | (Borowiec, 2016) |
| *Simopone* | 39 | ? | No | ants | ? | ? | ? | ? | ? | (Bolton and Fisher, 2012; Borowiec, 2016) |
| *Tanipone* | 10 | ? | No | ? | ? | ? | ? | ? | ? | (Borowiec, 2016) |
| *Vicinopone* | 1 | ? | No | ? | ? | ? | ? | ? | ? | (Borowiec, 2016) |
| *Parasyscia* | 51 | 100 | No | ants | ? | ? | few | no | group | (Brown, 1975; Ito et al., 2018; Wilson, 1958a) |
| *Zasphinctus* | 23 | 100 | No | ants | ? | inside nest (inferred) | few | no | group | (Briese, 1984; Buschinger et al., 1989; Clark, 1923; Wilson, 1958a) |
| *Lividopone* | 1 | 10 | No | ants | ? | ? | few | no | group | (Borowiec, 2016; Brown, 1975) |
| *Lioponera* | 74 | 10 | No | ants | scout-initiated | inside nest | few | no | group | (Brown, 1975; Clark, 1923, 1941; Hölldobler, 1982; Wheeler, 1918; Wilson, 1958a) |

Table S2: Classification (from the literature) of maximum colony size (order of magnitude), prey spectrum, foraging characteristics, and overall type of foraging behavior for each extant doryline genus (see Methods for details of classification). We also list the number of described species for each genus, and whether or not they are classified as army ants.
(*) The classification of *Aenictogiton* as an army ant is currently based solely on its phylogenetic position; colonies of this genus have not yet been observed.

| | Colony size | | Raiding behavior | | Prey spectrum | |
|---|---|---|---|---|---|---|
| Doryline MRCA | **Small (<5 * $10^4$)** | 0.994 | **Group raiding** | 0.93 | **Ants** | 0.991 |
| | Big (>5 * $10^4$) | 0.006 | Mass raiding | 0.07 | Termites | 0.004 |
| | | | | | Broad | 0.004 |

Table S3: Proportional likelihoods of each character state for the most recent common ancestor (MRCA) of dorylines from a one-parameter maximum likelihood (ML) reconstruction. Most parsimonious states from maximum parsimony (MP) reconstructions are in bold font.

| | | Initial recruitment location | | | | | | | | |
|---|---|---|---|---|---|---|---|---|---|---|
| **Event** | **Colony** | **First recruitment** | **Second recruitment** | **Third recruitment** | **Fourth recruitment** | **Fifth recruitment** | **Sixth recruitment** | **Seventh recruitment** | **Number of ants in raid** | **Bifurcations?** |
| 1 | 1 | outside | outside | outside | outside | outside | outside | NA | >1000 | Yes |
| 2 | 1 | inside | inside | outside | outside | NA | NA | NA | >1000 | Yes |
| 3 | 1 | outside | outside | outside | NA | NA | NA | NA | >1000 | Yes |
| 4 | 1 | outside | outside | outside | outside | outside | outside | NA | >1000 | Yes |
| 5 | 1 | inside | outside | outside | NA | NA | NA | NA | >1000 | Yes |
| 6 | 1 | inside | outside | outside | NA | NA | NA | NA | >1000 | Yes |
| 7 | 1 | outside | outside | outside | outside | outside | outside | NA | >1000 | Yes |
| 8 | 2 | outside | outside | outside | outside | outside | NA | NA | >1000 | Yes |
| 9 | 2 | outside | outside | outside | outside | outside | outside | outside | >1000 | Yes |
| 10 | 2 | outside | outside | outside | outside | NA | NA | NA | >1000 | Yes |

Table S4: Manually annotated raids and their features from two *O. biroi* colonies with ca. 5,000 workers each.

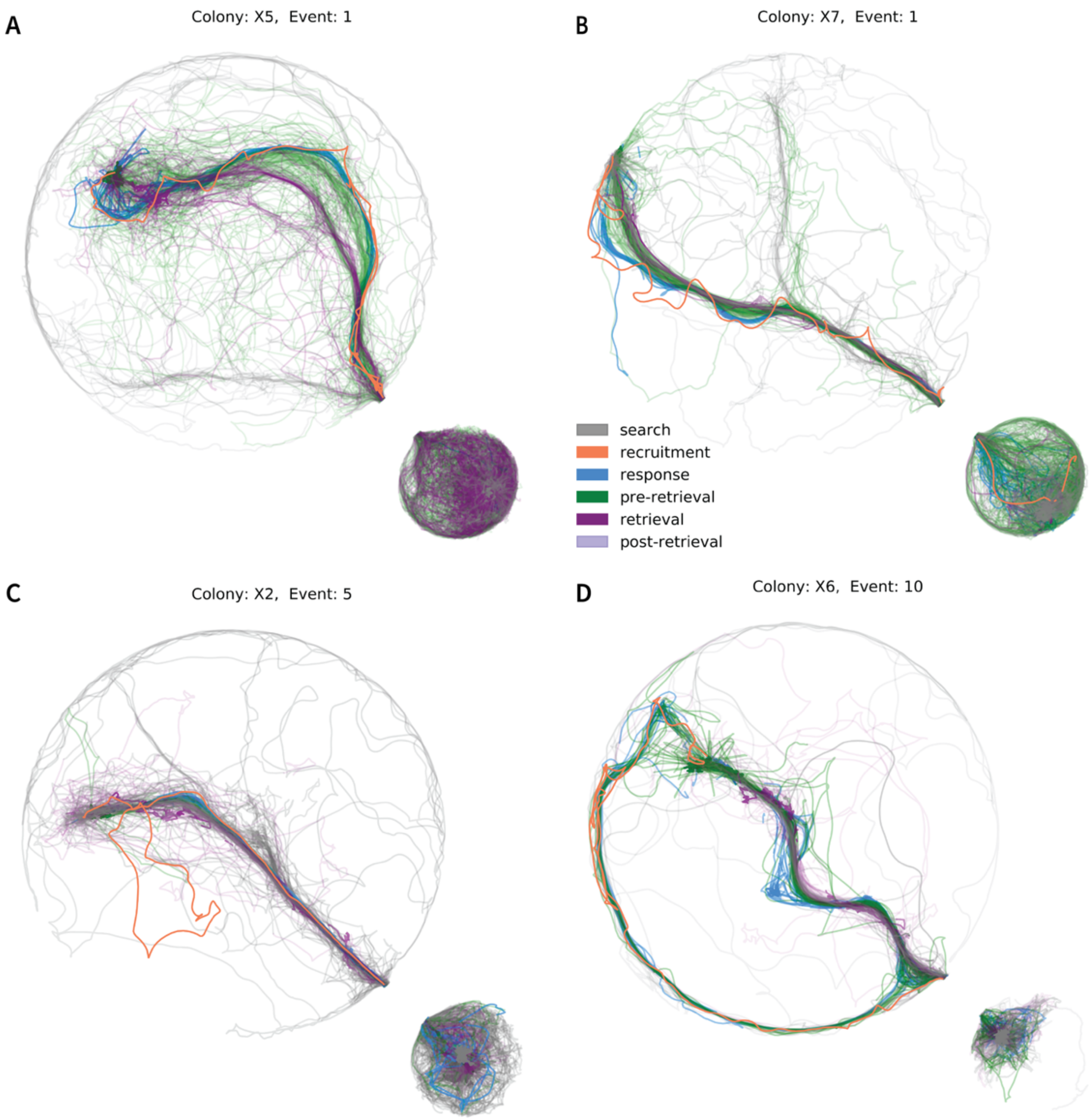


Figure S1: Additional examples of raids (Movies S3 to S6), showing that the coarse structure of the raid is similar across events, despite differences in details. For instance, in (A), the trail shortens discretely during the pre-retrieval phase (with continuous trail shortening in this phase evident in all events). In (C), the scout's trajectory homewards is somewhat meandering, looping over itself. However, the responding ants ignore this loop. In (D), the response trail bifurcates, with most ants not following their recruiter's path to the food. Careful observation of this event (Movie S6) suggests that two ants laid trail pheromone from the food to the nest simultaneously. The scout is defined as the ant that actually initiates the response, i.e., the first ant to enter the nest. When the ants leave the nest in response to her recruitment, they encounter the second recruiting ant near the nest entrance, who appears to have laid the trail that they then largely follow to the food. This event illustrates that colonies of 25 ants may have multiple scouts that do not appear to coordinate their behavior.

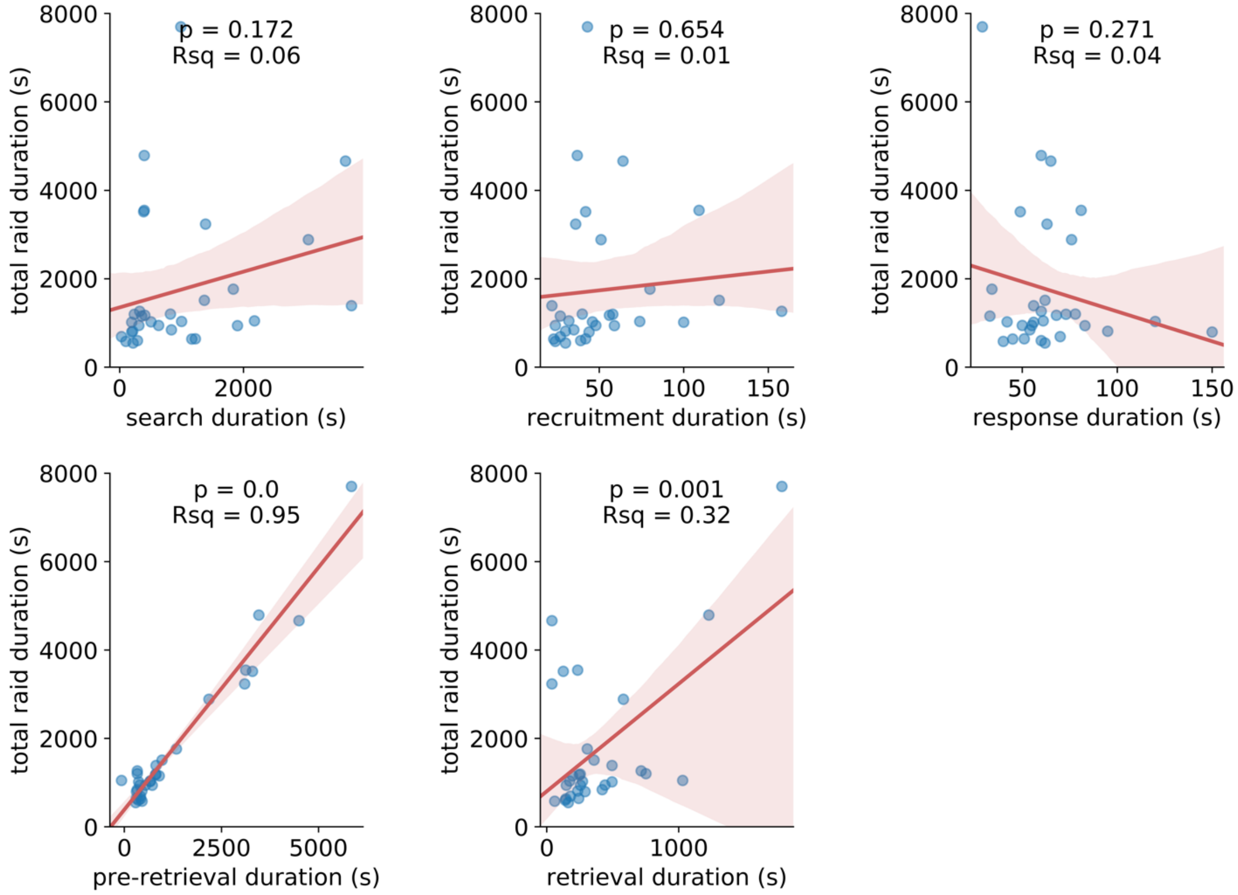


Figure S2: Variation in the duration of the ‘pre-retrieval’ phase explains most variation in the duration of raids (see also Figure 1D), while variation in the duration of the retrieval phase may also explain some variation in total raid length. Red lines depict the linear regression lines-of-best-fit, and error bands depict the 95% CIs for each regression.

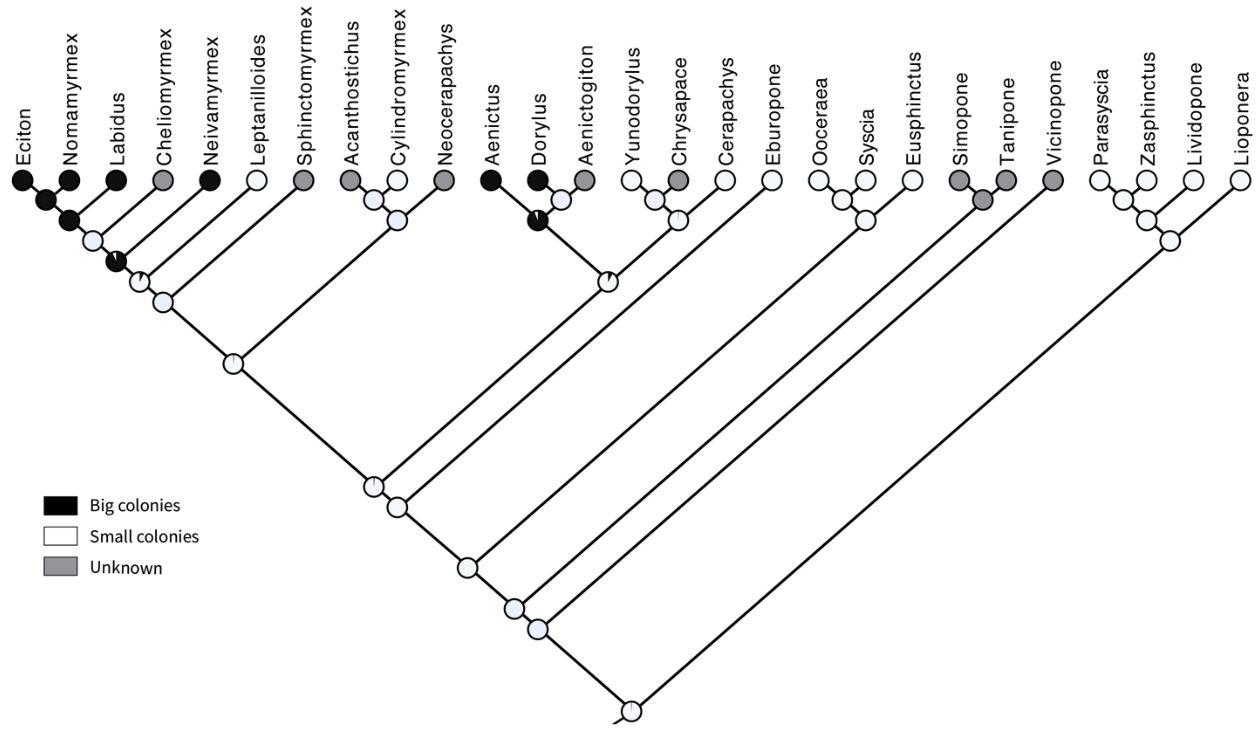


Figure S3: ML ancestral character state reconstruction for colony size. Pie charts at each node of the phylogeny depict the proportional likelihoods of both possible colony size states. The doryline MRCA (at the base of the tree) is highly likely to have had small colonies. As described in the Supplementary Methods, genus-level colony size states were binarized, with colonies above a threshold of $5*10^4$ workers classified as big, while all other colonies were classified as small. See Table S3 for proportional likelihoods and MP reconstructions.

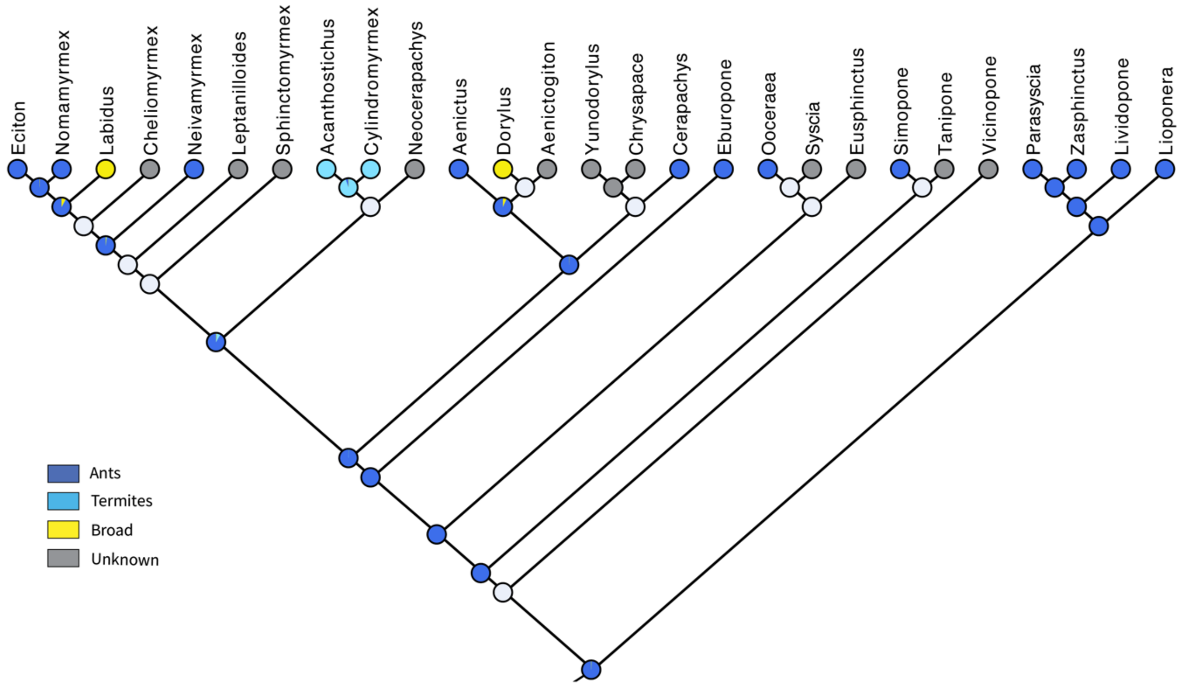


Figure S4: ML ancestral character state reconstruction for prey spectrum. Pie charts at each node of the phylogeny depict the proportional likelihoods of each possible prey spectrum state. The doryline MRCA (at the base of the tree) is highly likely to have been myrmecophagous (i.e., an ant-predator). See Table S3 for proportional likelihoods and MP reconstructions.

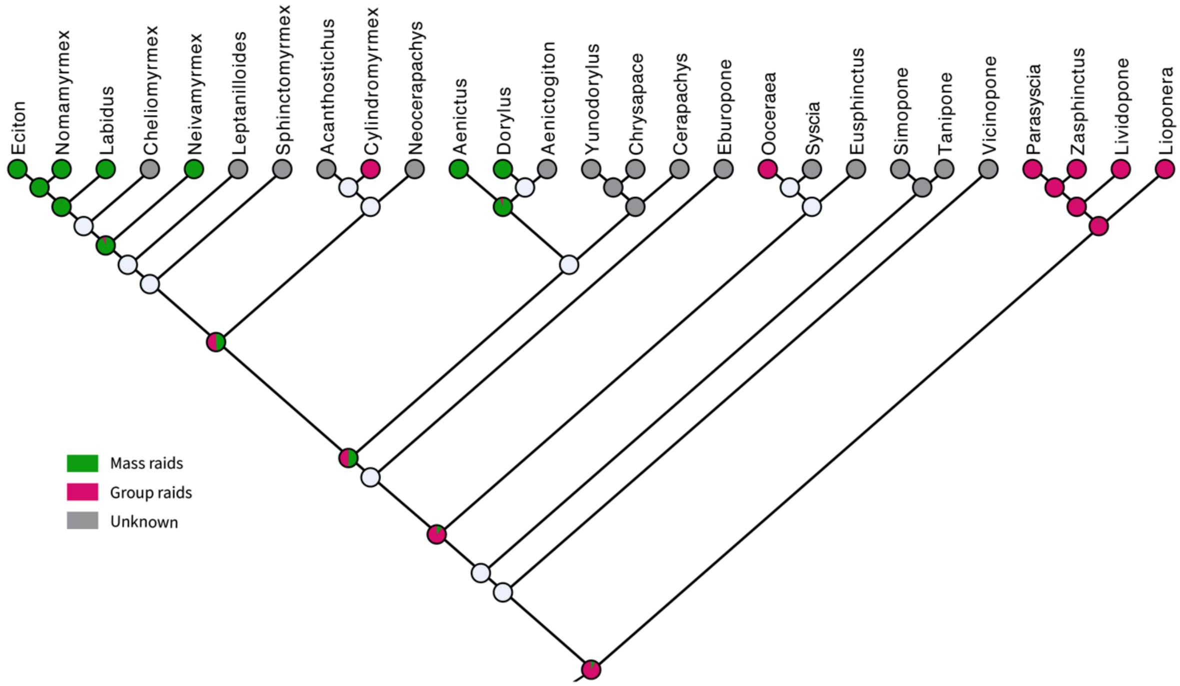


Figure S5: ML ancestral character state reconstruction for raiding behavior. Pie charts at each node of the phylogeny depict the proportional likelihoods of both possible raiding behavior states. The doryline MRCA (at the base of the tree) is highly likely to have been a group raider. See Table S3 for proportional likelihoods and MP reconstructions.

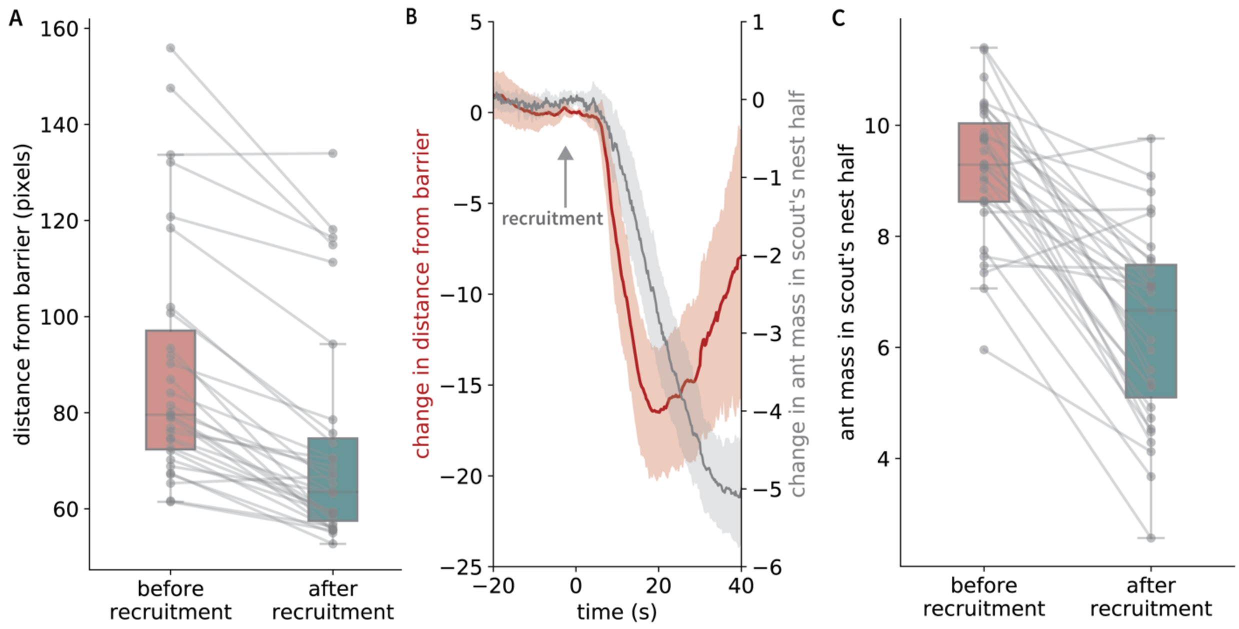


Figure S6: Additional analyses for the experiment in Figure 3 A and B, showing that in a modified arena with a porous barrier down the middle of the nest, ants on the scout's side leave their nest half soon after recruitment, while ants on the other side run towards the barrier. (A) A Wilcoxon test shows that the center of mass of the ants (on the side of the nest that does not have food) is closer to the barrier separating the two nest halves after recruitment compared to before recruitment ($n = 31$ events, $p=1.5*10^{-6}$), demonstrating that the recruitment pheromone is attractive. (B) The timecourse of change in the ant mass (i.e., the normalized number of dark pixels – a proxy for the number of ants; see Methods) in the scout's nest half in gray, overlaid in red with the timecourse of the change in distance from the barrier of the ants in the other nest half. All time courses are aligned to the time of recruitment of their respective events. The dark lines represent the means of the timecourses, and the error bands depict the 95% confidence intervals of the means. (C) A Wilcoxon test shows that the ant mass in the scout's nest half decreases after recruitment ($n = 31$, $p=8.9*10^{-7}$). Boxes represent the interquartile range and whiskers represent 1.5 times the interquartile range in panels (A) and (C), and the gray lines connect paired datapoints representing the same colony before and after recruitment during the same event.

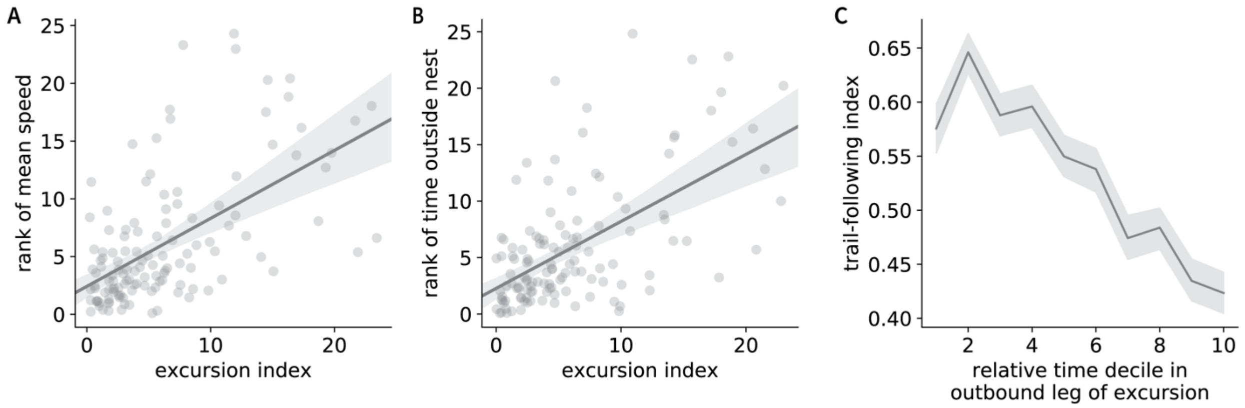


Figure S7: Analysis of individual ants' excursions in the search phase. (A) Ants typically walk faster in later excursions (n = 127 excursions, linear regression $p=4.5*10^{-13}$), and (B) spend longer outside (n = 127 excursions, linear regression $p=3.1*10^{-13}$). (C) Each excursion typically begins with a period of trail-following, followed by a period of traversing untrodden ground before reversal, as suggested by comparison of a trail-following index constructed from a binary heatmap classification (see Methods) against the relative time decile into the outbound leg of each excursion. Here, the dark line connects the means of each decile, while in (A) and (B) it represents the regression line. The error band depicts the 95% confidence interval of the mean in (C), and the 95% confidence interval of the regression line in (A) and (B).

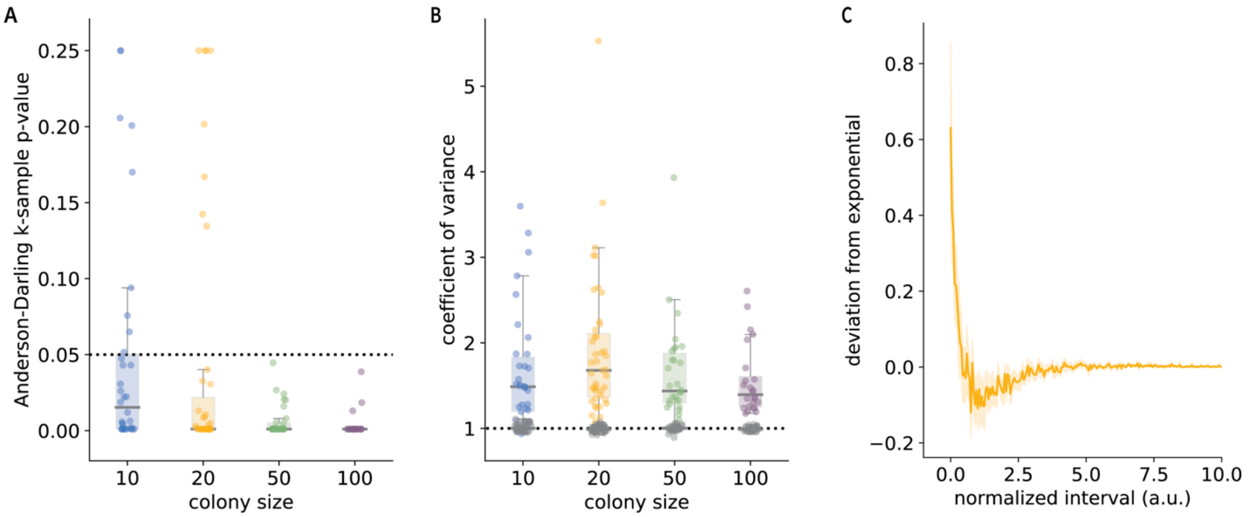


Figure S8: Inter-exit interval distributions across all colony sizes typically deviate significantly from the random, exponential expectation in the same way, with an overrepresentation of short intervals. (A) Distributions of p-values from Anderson-Darling k-sample tests comparing all real inter-exit interval distributions from all colonies to matched, simulated exponential distributions shows that across all tested colony sizes, a majority of interval distributions are unlikely to be explained by a random, Poisson process (n ≥ 24 events for each colony size). The majority of interval distributions lie below the significance threshold of p=0.05 (depicted as a black dotted line). (B) Distributions of coefficients of variance show that across all colony sizes the value is higher than would be expected for an exponential distribution (this expectation of one is depicted as a black dotted line, and gray data points show coefficients of variance for simulated exponential distributions). This shows that for all colony sizes short exit intervals are overrepresented. Boxes represent the interquartile range and whiskers represent 1.5 times the interquartile range in both panels. (C) Ants leave the nest in close proximity to each other more often than expected by chance. The y-axis shows that on average the deviations between real distributions and matched, simulated exponential distributions are greatest in the smallest time bins for colonies of size 20. The x-axis represents the normalized intervals between subsequent nest exits (note that the units are arbitrary as the mean exit interval for each sequence of inter-exit intervals is set to a value of one). The dark amber line shows the mean across all sequences, and the error band represents the 95% confidence interval of the mean.

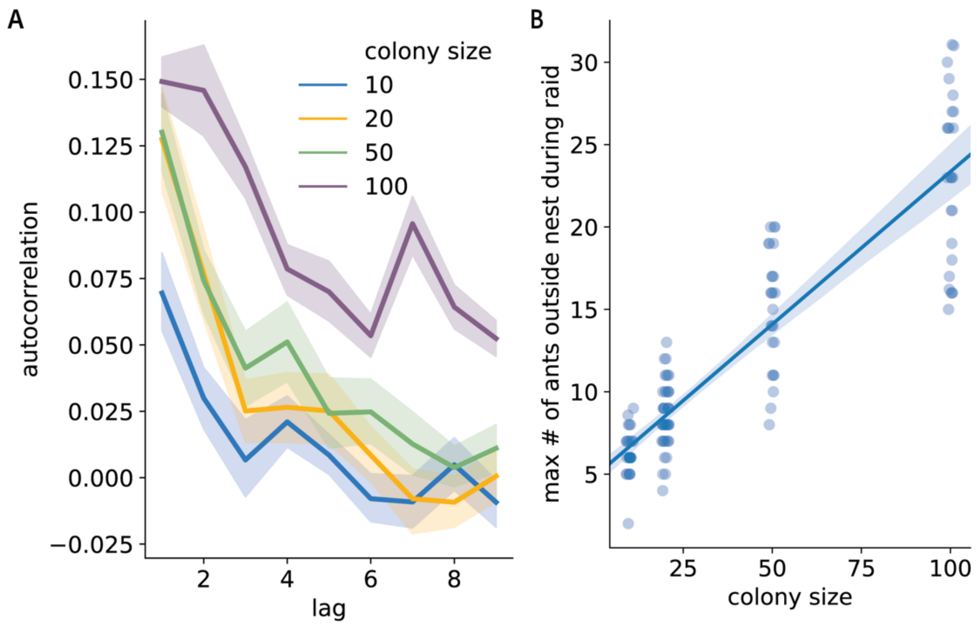


Figure S9: As colony size increases, the autocorrelation of spontaneous inter-exit intervals also increases, and so does the number of ants that participate in raids. (A) The magnitude and lag of the autocorrelation increases with colony size. Dark lines depict mean autocorrelation values for the detrended sequence of inter-exit interval sequences across colony size, and error bands depict standard error of the mean (n ≥ 24 for each colony size). (B) The estimated number of ants that participate in raids increases as a function of colony size. The y-axis depicts an estimate of the maximum number of ants outside the nest during raids; the x-axis values are jittered to aid visualization (n = 126 raids, linear regression p<0.0001).

Movie S1: This video depicts a representative group raid in a colony of 25 individually tagged *O. biroi* workers. The nest is at the bottom right (small circle), and the foraging arena is at the top left (large circle). The food – a single fire ant pupa, dyed blue – is visible in the foraging arena. The end of the search phase, and the entire recruitment and response phase, can be seen. This video shows the raiding event represented in Figure 1 A and B and is sped up 3x. For presentation purposes, the video was processed by masking the area outside the raiding arena, the nest and the connecting tunnel, and background-subtracting the region inside the foraging arena to eliminate reflections and distractors such as trash. Individual ant identities and recent tracks (i.e., the last five seconds) were superimposed. The time counter at the bottom left depicts time (in hours:minutes:seconds) since the start of video recording. Unprocessed videos are available upon request from the authors.

Movie S2: This video shows the same raid as Movie S1. The video includes the entire raid, from the beginning of the search phase to the end of the post-retrieval phase, for the raiding event described in Figure 1, A and B. The video is sped up 10x. The video was processed as described in the caption of Movie S1.

Movie S3 – S6: These videos show four additional raiding events from the experiment involving colonies of 25 individually tagged *O. biroi* workers. The food – a single fire ant pupa, dyed blue – is visible in the foraging arena. See Figure S1 and its associated legend for images of the ants' tracks, as well as descriptions of idiosyncrasies of each raid shown here. These videos show that, despite variation in specific aspects of the raid, the coarse spatial and temporal structure of group raids is stereotyped. All videos are sped up 3x. Videos were processed as described in the caption of Movie S1.

Movie S7: This video shows a representative recruitment and response phase from a modified arena with a barrier in the middle of the nest. A single fire ant pupa is placed at the top of the foraging arena connected to the left nest half. A scout locates the food and recruits ants from her nest half to it, while the ants in the other nest half respond to her recruitment by running into the barrier separating the nest halves. See also Figure 2, A and B, and Figure S6. The video is sped up 3x. For presentation purposes, the video was processed by masking the area outside the raiding arena and background-subtracting the region inside the foraging arena to eliminate reflections and distractors such as trash. As the food is not dyed, it is largely obscured during background subtraction, and is marked in the first few seconds of the video with a red circle. The time counter at the bottom left depicts time (in hours:minutes:seconds) since the start of video recording.

Movie S8: This video shows four representative group raids from colonies ranging in size from 10 to 100. Only recruitment and response phases from each raid are shown, to demonstrate that the coarse dynamics of the raid do not change with colony size. Videos are sped up 3x, and were processed as described in the caption of Movie S7.

Movie S9: This video shows a colony of 50 untagged *O. biroi* workers in the search phase of a raiding event. The food is at the top left (and is initially highlighted with a red circle). The ants have never left the nest before during this raiding event, i.e., at the beginning of the video, the foraging arena is void of trail pheromone. Ants can be seen spontaneously leaving the nest in serial excursions that build into a small column of workers that travels gradually away from the nest. The video is sped up 10x. The video was processed as described in the caption of Movie S7.

Movie S10: This video shows a representative recruitment event during a raid in a colony of ca. 5,000 *O. biroi* workers. The scout's position is briefly highlighted in red before she begins her recruitment run. The response to recruitment and trail bifurcation are clearly visible. The video is sped up 15x.
